# Exploring chromatographic dimensions for state-of-the-art proteomics applications

**DOI:** 10.1101/2025.08.16.670650

**Authors:** Alicia-Sophie Schebesta, Kathrin Korff, Ericka C.M. Itang, Vincent Albrecht, Philipp E. Geyer, Johannes B. Mueller-Reif

## Abstract

The evolution of mass spectrometry (MS)-based proteomics has been driven by continuous technological advances in sample preparation, instrumentation, and data acquisition. While chromatographic separation has historically been considered a critical bottleneck in achieving comprehensive proteome coverage, recent developments in ultra-fast data acquisition fundamentally challenge this paradigm. We investigated whether the traditional paradigm that chromatographic performance directly correlates with proteome depth still holds true. Spanning a matrix of experiments with five distinct stationary phases, including C18 chemistries, C8, and Phenyl-Hexyl, across eight column lengths (40-140 mm), we evaluate protein identification performance using data-independent acquisition (DIA) on the Orbitrap Astral mass spectrometer. Despite substantial chromatographic differences, we observed remarkably convergent proteome coverage metrics. All C18 and C8 phases consistently achieved over 150,000 precursor- and approximately 9,000 protein group identifications, regardless of column length variations. While distinct selectivity fingerprints persisted across chemistries, these chromatographic differences did not translate into meaningful variations in proteome coverage under high-speed acquisition conditions at 200 Hz. We conclude that the analytical bottleneck has fundamentally shifted from chromatographic resolution to mass spectrometric sampling efficiency, where comprehensive peptide identification is now gained through advanced spectral deconvolution rather than physical separation alone. This paradigmatic shift is reflected in modern proteomics by method development priorities being directed beyond traditional separation optimization, with greater emphasis placed on operational robustness, analytical throughput, and reproducibility.

## Introduction

Liquid chromatography coupled with mass spectrometry (LC-MS/MS) is the central analytical platform for proteome analysis, with reversed-phase high-performance liquid chromatography (RP-HPLC) widely used for peptide separation^1–6^. Traditional proteomics workflows have long relied on capillary columns, typically 75 to 150 μm inner diameter, 15 to 50 cm length, and sub-2 μm packing material, which have been operated with capillary to nanoflow systems to maximize sensitivity. This general setup has remained largely consistent for nearly two decades, even as mass spectrometer performance has advanced constantly.

The principles of reversed-phase chromatography imply that several parameters influence separation quality, including column chemistry, length, gradient conditions, and flow rate. Different stationary phases, such as C8 and C18 alkyl chains, polar-embedded groups, or Phenyl-Hexyl ligands, provide varying retention mechanisms, including hydrophobic interactions, hydrogen bonding, or π-π interactions^7,8^. These differences can affect peptide selectivity, particularly for analytes with challenging physicochemical properties. C18 phases typically provide the strongest hydrophobic retention and broadest applicability for peptide separations, while C8 phases offer reduced hydrophobic character that can benefit the analysis of highly hydrophobic peptides by reducing excessive retention and peak broadening. Phenyl-Hexyl phases introduce aromatic selectivity through π-π interactions with aromatic amino acid residues, potentially offering complementary separation mechanisms for peptides rich in phenylalanine, tyrosine, or tryptophan. Additionally, differences in silica particle morphology, pore structure, and surface area can significantly influence mass transfer kinetics and peak shape, with newer particle technologies designed to minimize band broadening and improve efficiency^9–11^. Similarly, increasing column length should enhance peak capacity through improved resolution^12,13^. However, this often comes at the cost of increased backpressure, extended runtimes, and diminishing gains under long gradient conditions.

Historically, the limited acquisition speed of early-generation mass spectrometers made chromatography a key factor in precursor identification performance. With instruments acquiring only ∼1 MS/MS scan per second, chromatographic peak shape, width, and resolution were critical to maximizing identifications^12^. Under these constraints, careful optimization of column material, length, and particle size was essential^14^.

In recent years, advances in MS instrumentation, such as the Orbitrap Astral with MS/MS rates of 200 Hz, have markedly increased acquisition speed and sensitivity^15–17^. Modern platforms can routinely quantify over 10,000 proteins in 30 minute runs and ∼7,000 proteins in just 5 minutes using data-independent acquisition (DIA)^16^. This has enabled immense advances in terms of gradient length required for deep proteome analysis. While MS-based proteomics measurements were mostly done in hour-long gradients only years ago, the increase in sensitivity and speed has made this practice mostly obsolete. The routine use of gradient runtimes on the minute scale has revolutionized the proteomics field in areas where high-throughput application is needed, like population proteomics, interaction profiling, drug target screening, and single-cell applications. These improvements were driven by advances in MS but also by LC innovations and software^18–20^. Here, we raise the question whether chromatographic optimization continues to have the same impact on identification rates in current high-throughput workflows. We hypothesize that the above-described technical advances may have reduced the relative impact of parameters such as peak capacity and resolution on overall identification performance in LC-MS-based proteomics experiments.

In this work, we systematically evaluate the impact of column material and length on LC-MS performance for state-of-the-art MS instrumentation under consistent conditions. We compare five different stationary phases, including C8, Phenyl-Hexyl, and two C18 materials (Exsil Mono 1.35 µm and Reprosil Saphir 1.3 µm and 1.5 µm), across column lengths ranging from 40 to 140 mm in 10- or 20-mm increments. Our aim was to assess whether differences in column chemistry and dimensions influence proteome coverage and, further, to determine where column selection retains value in shaping selectivity, including its relationship with chromatographic metrics such as peak width and resolution.

In summary, this study examines the extent to which column chemistry and length affect proteome coverage and selectivity in high-speed data-independent acquisition workflows, clarifying the relevance of traditional chromatographic optimization for current proteomics applications.

## Results

To benchmark the performances of various columns, we measured a tryptic digest of HeLa in triplicates. We span an experimental matrix of five different column chemistries (Exsil Mono C18 (1.35 µm; EM), Reprosil Saphir C18 (1.3 µm; RS13), Reprosil Saphir C18 (1.5 µm; RS15), Reprospher C8 (1.8 µm; C8), and Reprospher Phenyl-Hexyl (1.8 µm; PH)) and eight column lengths (40, 50, 60, 70, 80, 100, 120, 140 mm) (**Figure 1A**). Comparison of identified protein groups (**Figure 1B**) and precursors (**Figure 1C**) reveals broadly consistent performance across all conditions. The EM columns yield the highest identifications with an average of 9,200 protein groups and over 160,000 precursors, followed closely by RS13 and RS15 and C8, each exceeding 150,000 precursors and ∼9,100 protein groups. In contrast, PH consistently identifies the lowest number of proteins (∼8,400) and precursors (∼110,000).

**Figure 1:**
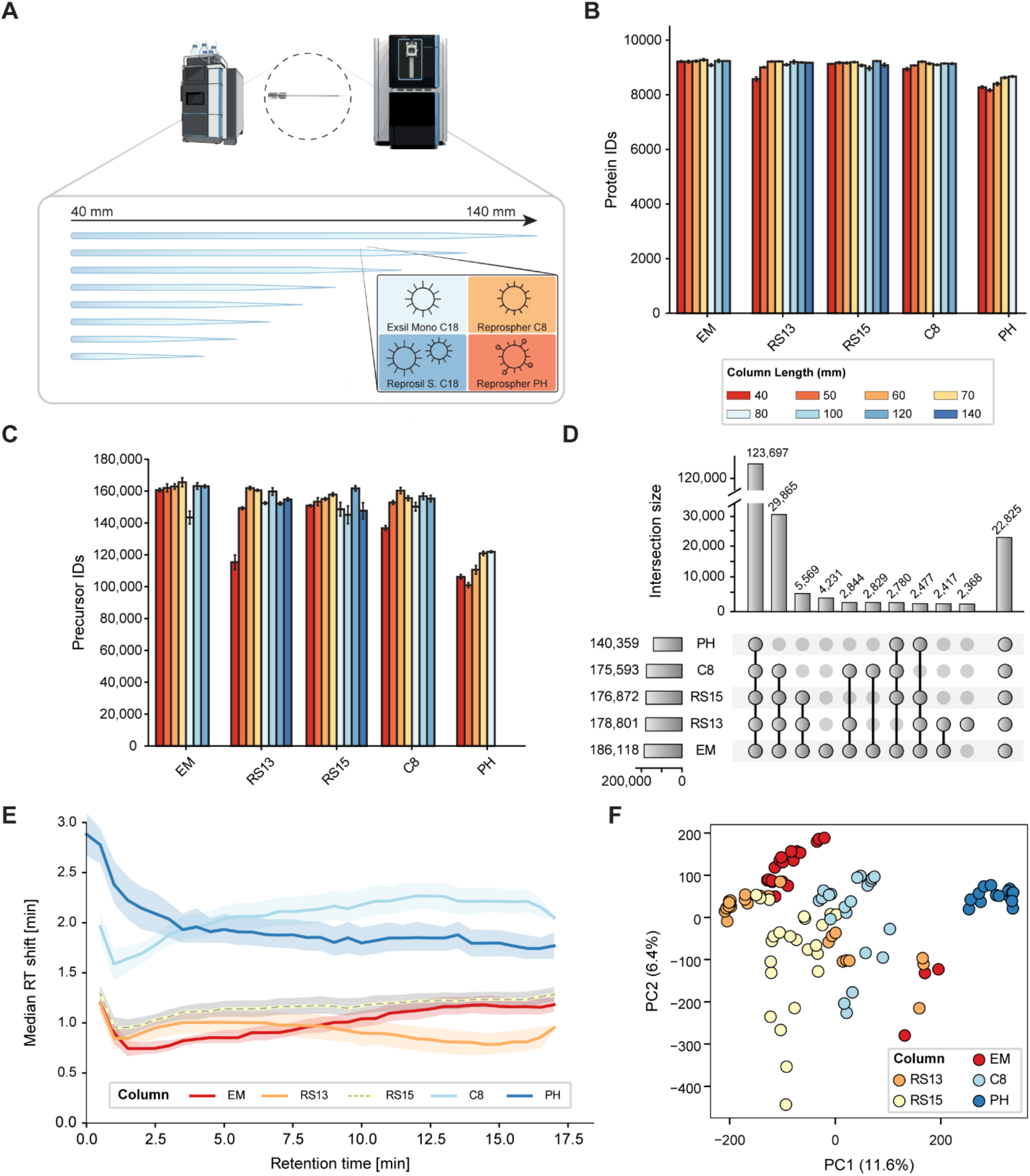
Column performance comparison across chemistries and lengths. (A) Schematic overview of experimental design testing five stationary phases across eight lengths. (B) Protein group identifications across column materials and lengths. (C) Precursor identifications across column materials and lengths. (D) UpSet plot showing precursor overlap between column chemistries at 70 mm. Only the top intersections are shown; all remaining overlaps and unique sets are aggregated in the final bar. (E) Median retention time differences (ΔRT = RT_80_ – RT_40_) plotted as a function of retention time from 40 mm columns. (F) PCA of precursor intensities colored by column chemistry. Missing values were imputed using k-nearest neighbor imputation (k = 5). Intensities were standardized (mean = 0, variance = 1) prior to principal component analysis.

While protein identifications remain largely unaffected by column length, precursor identifications show modest variation. The EM column demonstrates high performance across lengths, with only the 80 mm variant showing slightly lower performance. For RS13, identifications increase moderately from 40 to 60 mm before plateauing, whereas RS15 does not exhibit this trend. C8 displays gradually increasing precursor identifications with column length while maintaining stable protein-level performance. Remarkably, C8 achieves identification levels comparable to C18 chemistries despite its lower hydrophobicity, highlighting the capability of modern mass spectrometers to minimize separation-dependent losses.

These identification patterns are even more obvious when examining elution behavior over time. Plotting cumulative identifications across the gradient (**Supplementary Figure 1**) shows that over 80% of proteins are identified within the first third of the chromatographic run, with systematic shifts toward later elution as column length increases for all chemistries except PH. At the precursor level, the cumulative curves rise more gradually and plateau after approximately 15 minutes, reflecting the expected separation profile.

Given the similar identification performance across columns, we investigated whether the same precursors are being detected by examining overlap patterns using 70 mm column data (**Figure 1D**). More than 120,000 precursors (∼70%) are shared across all five columns, including the majority identified on PH. Additional overlap sets emerge between C8 and the C18 columns, and among the three C18 variants alone. Despite this extensive sharing, substantial unique identifications occur for EM and RS13 with 4,231 and 2,368 unique precursors, respectively. Similar patterns appear at the protein level (**Supplementary Figure 2**), with 8,833 proteins shared across all chemistries.

To quantify how column length affects retention across chemistries, we compared retention time differences between 40- and 80-mm columns **(Figure 1E**). PH and C8 show substantially larger shifts (∼1.7–2.3 min), roughly two-fold higher than C18 chemistries (∼0.8–1.3 min). Chemistry-specific patterns emerge across the gradient. PH exhibits maximum early shifts (∼2 min) that decrease toward late elution, while C8 shows the opposite, with increasing shifts exceeding 2 minutes late in the run. Among C18 phases, EM demonstrates minimal early sensitivity to column length (<0.7 min) that increases substantially to >1 min at late retention times, indicating precursor retention becomes progressively more affected by column length. RS13 shows the inverse behavior with ∼1 min early differences that decrease over the gradient, meaning column length impact diminishes for late-eluting precursors. RS15 remains nearly constant (∼1.0–1.2 min) throughout. These findings reveal that length-dependent retention varies dramatically by chemistry and gradient position. Supplementary Figure 3 provides distributional analysis across retention windows, while Supplementary Figure 4 shows individual GAPDH precursor chromatograms.

These chromatographic differences translate into distinct quantification profiles when examined globally through principal component analysis on precursor intensities (**Figure 1F, Supplementary Figure 5C,D**). Measurements cluster primarily by column chemistry, with PH forming a distinct group separated along PC1 (11.6% variance). EM and RS13 cluster closely together and separate from RS15 and C8 along PC2 (6.4% variance). This separation becomes even more pronounced at the protein level (**Supplementary Figure 5A,B**), where PH shows sharp separation along PC1 while C18 and C8 columns cluster closely together.

In summary, while identification depth remains remarkably consistent across column types and lengths, the underlying separation characteristics reveal meaningful differences. Column chemistry and length systematically influence elution profiles of shared precursors, with PH showing the most distinct behavior through altered retention patterns and broader peak shapes, while other columns, particularly EM, maintain stable performance and high reproducibility.

While Figure 1 demonstrated that most precursors are identified across all packing materials and column lengths, we evaluated chromatographic parameters to explore subtle differences in separation behavior. We analyzed full width at half maximum (FWHM) as a measure of peak sharpness and retention time correlations for separation characteristics.

FWHM distributions across column chemistries (**Figure 2A**) reveal that EM exhibits the sharpest peaks with a median FWHM of 0.050 min, followed closely by RS13 and RS15 at 0.054 and 0.057 min, respectively. These correspond to approximately 3 seconds in width at half maximum. C8 and PH columns show broader elution with median FWHMs of 0.060 and 0.100 min, respectively. Column length has minimal impact on peak width for most chemistries, with the three best-performing columns showing nearly identical FWHM distributions across all lengths. C8 demonstrates more pronounced length dependence, with peaks becoming 13% sharper from 40 mm to 120 mm columns. PH exhibits the broadest peaks overall, lacking distinct modal distributions and showing only slight improvement with increased length.

**Figure 2:**
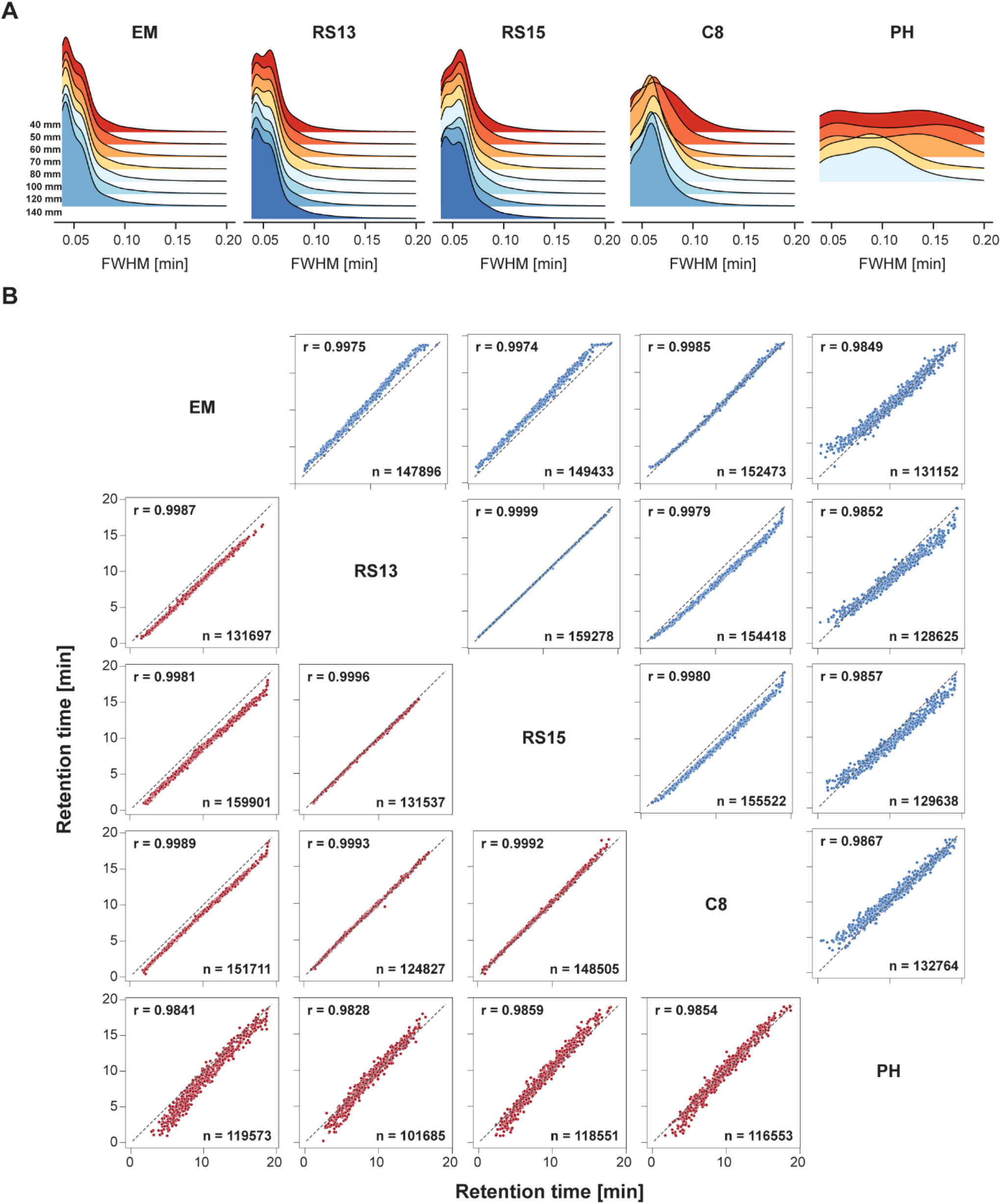
Chromatographic performance comparison across column chemistries and lengths. (A) FWHM distributions for each column type across different column lengths (40-140mm). Data only shown from 0 to 0.2 FWHM. (B) Retention time correlation matrix for 40mm (red) and 80mm (blue) columns showing pairwise comparisons between column types. Only 0.5% of all data points are shown.

Retention time correlation analysis between shared precursors reveals high reproducibility across most column types **(Figure 2B**). Correlations between EM and both RS variants are near perfect (r >0.98), with precursors eluting slightly earlier on RS columns. This shift increases modestly with retention time but remains consistent and predictable.

C8 correlations are similarly strong, particularly for 80 mm columns, where retention times closely match EM, though late-eluting hydrophobic precursors show earlier elution on C8 compared to C18 chemistries, reflecting the expected lower retention capacity. PH demonstrates distinct behavior with slightly lower correlations (r = 0.98-0.99) and broader scatter patterns. Early-eluting precursors that appear at ∼5 minutes on other columns elute around minute 2 on PH, indicating reduced retention of hydrophilic precursors. Conversely, hydrophobic precursors show enhanced retention on PH compared to other columns, creating a shifted separation profile with different selectivity characteristics.

Analysis across column lengths within individual chemistries shows near-perfect correlations (r > 0.99) for both RS13 and EM (**Supplementary Figures 6, 7**). Longer columns shift elution to later times in the gradient due to increased retention capacity, with this effect being more pronounced for EM, suggesting greater peptide length dependence for this chemistry.

These findings demonstrate that while overall performance remains comparable, column chemistry introduces systematic differences in peak sharpness and retention selectivity. These variations, particularly for PH and C8 chemistries, may be leveraged for enhanced separation of specific precursor populations with distinct physicochemical properties.

While the majority of precursors overlap across all five column chemistries (**Figure 1B**), a subset of precursors is uniquely identified by individual columns, warranting detailed investigation (**Figure 3A**). From a total of 201,702 identified precursors, 5% are detected exclusively in a single column type, with the EM column accounting for 2.1% (4,231 precursors) and RS13 contributing 1.2% of unique identifications. To ensure reproducibility, we filtered unique precursors to include only those consistently identified across all three technical replicates. This stringent filtering yielded 1,278 of 4,231 precursors for EM (30%), 594 of 2,368 for RS13 (25%), and 1,131 of 1,915 for PH (59%). Notably, PH demonstrates the highest reproducibility among unique precursors, reflecting the distinct selectivity conferred by its alternative stationary phase chemistry.

**Figure 3:**
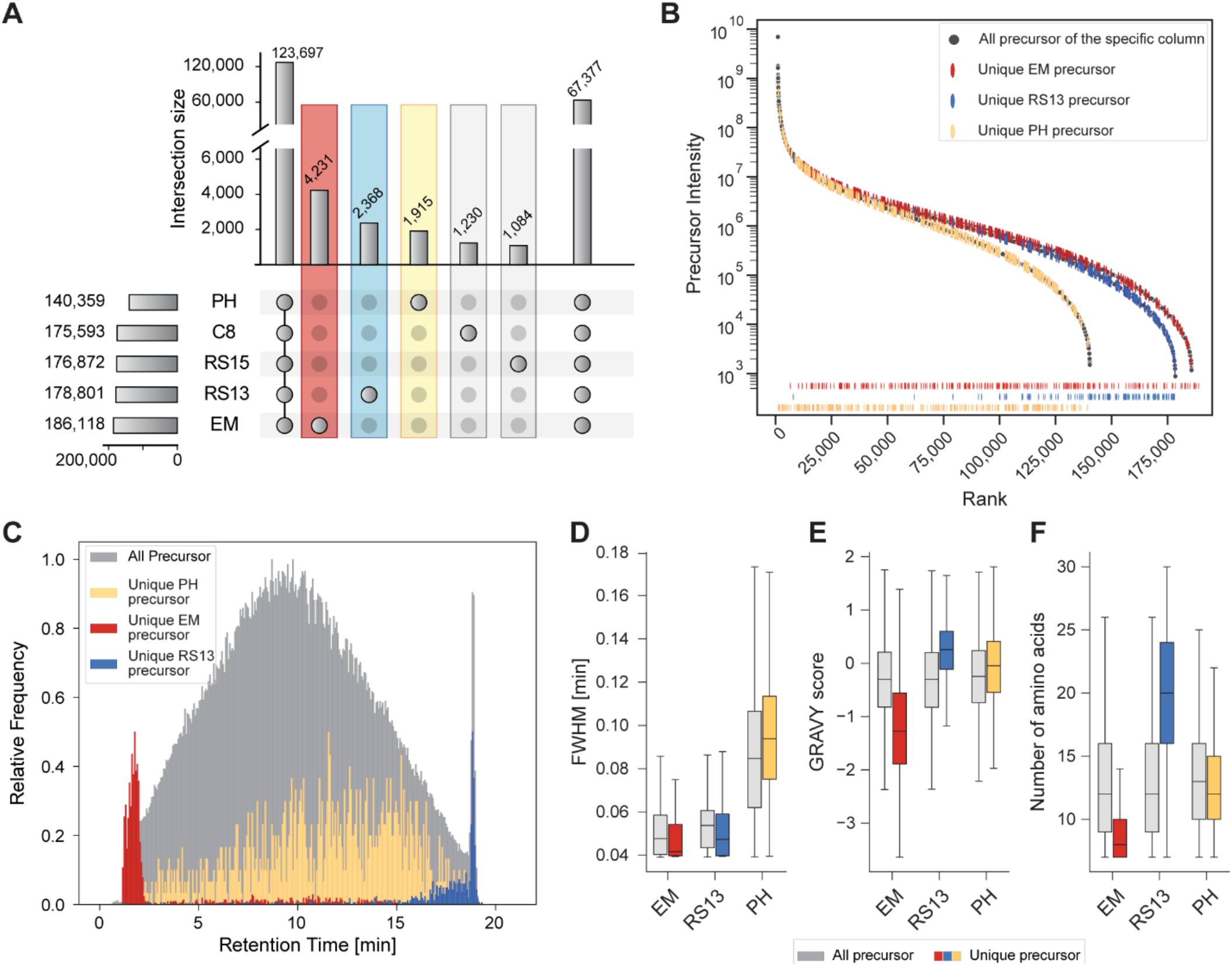
Unique precursor characterization across column chemistries. (A) Modified UpSet plot showing precursors unique to each column (colored bars) along with the number shared across all five columns (left) and the sum of all remaining intersections (right). Intermediate overlaps between columns are not shown individually. (B) Shared rank-abundance plot for three selected columns: EM (red), RS13 (blue), and PH (yellow). The lower panel shows the distribution of unique precursors along the rank axis for each column.; only 2% of all data points are shown. (C) Retention time distributions of all precursors (grey) and unique precursors (colored). Distribution of all precursors is normalized to 1 and distributions of the unique precursors is normalized to 0.5. (D) FWHM comparison between all (grey) and unique (colored) precursors. (E) GRAVY score distributions comparing hydrophobicity between all (grey) and unique (colored) precursors. (F) Amino acid length distributions comparing peptide length between all (grey) and unique (colored) precursors.

Given the substantial differences observed, we focused the following analyses on the three columns exhibiting the most pronounced selectivity differences. Rank abundance analysis (**Figure 3B, Supplementary Figure 8A**) reveals that unique precursors span the complete intensity range for EM and PH columns, demonstrating that these identifications are not merely low-abundance artifacts, but also include highly abundant species. In contrast, unique precursors from RS13 are predominantly ranked in the lower half of the abundance distribution.

Analysis of chromatographic elution profiles reveals distinct retention time signatures for unique precursors (**Figure 3C, Supplementary Figure 8B**). Unique EM precursors predominantly elute early in the gradient (1-3 minutes), while unique RS13 precursors elute exclusively at the gradient terminus (>15 minutes, with a pronounced peak at 19 minutes). This extreme elution behavior suggests that the bead size and material chemistry enable the separation of precursors that are not even identified by any other column type. In contrast, unique PH precursors exhibit a retention time distribution spanning the entire chromatographic range, though slightly shifted toward later elution compared to the overall precursor population. These distinct elution profiles indicate fundamental differences in physicochemical properties between unique precursor populations.

We therefore analyzed three key parameters to characterize these differences: full width at half maximum (FWHM) as a measure of peak width, GRAVY score as an indicator of hydrophobicity, and amino acid length. Peak width analysis (**Figure 3D**) shows that PH columns generate broader chromatographic peaks with approximately double the FWHM compared to other columns. For both EM and RS13, uniquely identified precursors exhibit narrower peak widths than the overall precursor population, despite their opposing elution profiles.

GRAVY score analysis (**Figure 3E**) demonstrates dramatic differences between unique precursor populations. While overall precursor populations show minimal differences across columns (median ≈ 0.3), unique EM precursors exhibit extreme hydrophilicity (median GRAVY = -1.3), reflecting their early elution and minimal retention by the C18 stationary phase. Conversely, unique RS13 precursors display strong hydrophobicity (median GRAVY ≈ +1.0), explaining their late elution and strong retention. Unique PH precursors show only slightly elevated hydrophobicity scores, consistent with their marginal retention time shift.

Precursor length analysis (**Figure 3F**) reveals comparable median lengths for overall precursor populations (12 amino acids for EM and RS13, 13 for PH). However, striking differences emerge for unique precursors. Unique EM precursors are substantially shorter (median 8 amino acids), representing a one-third reduction compared to the overall population. Conversely, unique RS13 precursors are remarkably longer (median 20 amino acids), representing a >60% increase with complete separation between the two distributions. PH shows only a marginal difference between unique and total precursor lengths.

These findings demonstrate that despite comparable overall performance metrics, different column chemistries provide distinct selectivity for precursor subsets with specific physicochemical properties. This selectivity proves particularly valuable for applications requiring enhanced detection of very hydrophilic or hydrophobic peptides, suggesting that column selection should be tailored to specific analytical requirements rather than relying solely on general performance criteria.

## Discussion

Our systematic evaluation of five column chemistries across eight lengths using state-of-the-art DIA-MS reveals a change in the role of chromatographic performance in contemporary LC-MS-based proteomics setups and workflows. Despite substantial differences in stationary phase chemistry and column length, overall proteome coverage was strikingly similar; all C18 and C8 phases yielded >150,000 precursors and ∼9,000 protein groups from HeLa digests. These performance metrics complement findings reported by Guzman et al. for nDIA methodology on human cell lysates^16^.

While we observed clear and consistent differences in selectivity and peak width between chemistries, these did not translate into substantial changes in overall identification depth under our acquisition conditions. This suggests that, in high-speed DIA workflows such as those enabled by the Orbitrap Astral MS, optimization of chromatographic performance has less impact on total proteome coverage than historically reported.

This finding appears to challenge aspects of established proteomics orthodoxy^1,2^. Traditional workflows, where chromatography constituted the primary analytical bottleneck, drove the development of ultra-high-pressure packing methods that could increase proteome depth by up to 35%^21^. At that time, even with faster MS instrumentation, it was argued that better chromatography remained essential to fully leverage instrument capabilities^22^.

Our results show that with modern DIA acquisition, this link between chromatographic quality and identification success is reduced, as >200 Hz acquisition rates enable comprehensive peptide identification through spectral deconvolution rather than chromatographic separation ^2,16,17^. It is important to note, however, that the extent of this effect is likely influenced by instrument capabilities and acquisition strategy, and may differ under other analytical configurations. To our knowledge, this is the first systematic evaluation under Astral DIA conditions, showing that analytical bottlenecks can shift from separation-limited to MS-limited performance, altering optimization priorities compared to those established over decades of method development^23,24^.

The existence of distinct selectivity patterns within this convergent performance landscape highlights how column chemistry may still play a role in future analytical strategies. Our analysis of orthogonal “selectivity fingerprints” shows that, although bulk identification metrics have converged, each chemistry still accesses complementary regions of physicochemical space. For example, EM preferentially identified early-eluting hydrophilic peptides, whereas RS favoured late-eluting hydrophobic sequences. This indicates that the two chemistries operate at opposite extremes of the retention window. PH showed a unique but more limited aromatic selectivity, likely driven by π–π interactions^7,8,25^. These differences suggest that column choice now serves more of a strategic than an essential role. Mechanistically, the contrast between monomeric and polymeric C18 bonding, and the resulting differences in surface uniformity, likely underlie these effects^9,26^. A similar shift in perspective applies to column dimensions. Classical chromatographic theory predicts substantial benefits from increased plate numbers, yet in our data, extending columns from 40 mm to 140 mm, and thereby reducing peak width, yielded only minimal identification gains^27–29^.

In our hands, these findings establish a new framework for proteomics method development that prioritizes analytical objectives over legacy optimization assumptions. For total proteome profiling, we document a performance equivalence across different column lengths and even packing phases, providing us with unprecedented flexibility based on practical rather than analytical constraints. However, the orthogonal selectivity fingerprints preserve opportunities for strategic optimization when targeting specific peptide populations, aligning with emerging multi-phase screening approaches^8,30^. With proteomics studies reaching sizes of tens of thousands of samples annually ^31,32^, and the minimal performance differences between column chemistries and lengths, the priority for selection of a chromatographic column is shifted towards reproducibility and operational lifetime, thus seeking robustness over long time periods. Consistent batch-to-batch performance and extended column longevity are now as critical as selectivity for minimizing variability and costs in sustained proteomics campaigns^6^, especially considering the increased throughput per day on a chromatographic system compared to only years ago. This fit-for-purpose philosophy represents a fundamental departure from the comprehensive optimization approaches that dominated separation-limited workflows^33,34^.

The limiting context of our findings is the application for full proteome analysis, acquired under standardized DIA conditions. Chromatographic optimization encompasses numerous variables; in this study, we focused specifically on column chemistry and length, while keeping other influential parameters such as gradient duration and flow rate constant. These factors may exhibit different optimization relationships with modern MS instrumentation and were beyond the scope of our current evaluation. Specific specimens, such as plasma or single cells, may exhibit different chromatographic dependencies due to distinct physicochemical compositions or dynamic range, while samples enriched for post-translational modifications^35^ or analyzed using DDA strategies may show stronger column dependencies than observed here^36^. These limitations define boundaries where traditional optimization principles may retain greater relevance, though our core findings likely extend broadly across contemporary proteomics applications.

The implications of our results extend beyond column selection and may help guide the redefinition of optimization priorities across proteomics workflows. We established the principle that MS capabilities can supersede traditional chromatographic constraints, suggesting that analytical bottlenecks in high-speed DIA workflows shift from chromatography to MS-centric considerations. In this setting, chromatographic optimization increasingly focuses on practical aspects such as reproducibility and column lifetime, supporting a more strategic rather than exhaustive approach to maximizing analytical success in next-generation proteomics platforms.

## Experimental procedures

### Experimental Design

#### HeLa cell cultivation

HeLa cells (S3 subclone, ATCC) were maintained in Dulbecco’s modified Eagle’s medium supplemented with 10% fetal bovine serum, 20 mM glutamine, and 1% penicillin-streptomycin at 37°C in a humidified 5% CO2 atmosphere. Mycoplasma testing was performed regularly to ensure culture sterility.

Cells were grown to approximately 80% confluence before harvesting. Following trypsinization with 0.25% trypsin/EDTA solution, cells were collected and transferred to 15 mL conical tubes. Cell pellets were washed twice using cold Tris-buffered saline (TBS) and recovered by centrifugation at 200g for 10 minutes. After removing the supernatant, cell pellets were immediately frozen in liquid nitrogen and stored at -80°C for subsequent analysis.

#### Cell lysate preparation

Cell pellets were lysed using the PREOMICS lysis buffer (PreOmics, Martinsired, Germany). The lysis protocol involved heating samples at 95°C for 15 minutes with continuous agitation at 1,500 rpm, followed by probe sonication using short pulses (10 cycles of 5 seconds on/off at 20% amplitude). Cellular debris was removed by high-speed centrifugation for 5 minutes.

Protein quantification was performed using tryptophan fluorescence measurement. For proteolytic digestion, both LysC and trypsin were added at a 1:100 enzyme-to-protein ratio (w/w) and incubated overnight at 37°C with gentle agitation at 1,500 rpm. The digestion reaction was terminated by acidification with 1% trifluoroacetic acid (TFA). Peptides were purified using styrene-divinylbenzene-reversed phase sulfonate (SDB-RPS) solid-phase extraction and reconstituted in 0.1% formic acid with 2% acetonitrile for LC-MS/MS analysis.

#### Data acquisition

All columns used in this study were 75 μm inner diameter pulled-tip capillaries supplied by Dr. Maisch GmbH (Ammerbuch, Germany). Five different stationary phases were systematically evaluated across eight column lengths (40, 50, 60, 70, 80, 100, 120, and 140 mm) to assess the impact of column chemistry and length on separation performance.

The following stationary phases and particle sizes were tested: Exsil Mono 100 C18 (1.35 μm particles, Cat. No. 6136973), Reprosil Saphir 100 C18 (1.3 μm particles, Cat. No. ra113.9e), Reprosil Saphir 100 C18 (1.5 μm particles, Cat. No. ra115.9e), Reprospher 100 C8 (1.8 μm particles, Cat. No. rs118.8e), and Reprospher 100 Phenyl-Hexyl (1.8 μm particles, Cat. No. rs118.ph). All columns featured integrated electrospray emitter tips created by laser pulling. No column oven was used; columns were operated at ambient laboratory temperature.

A 200 ng aliquot of HeLa tryptic digest was injected for each run, analyzed using a Thermo Scientific Vanquish Neo LC system coupled to an Orbitrap Astral mass spectrometer (Thermo Fisher Scientific) ^17,37^. The LC system was operated at a maximum pressure of 1500 bar with direct injection capability. Mobile phases consisted of 0.1% formic acid in water (buffer A) and 0.1% formic acid in acetonitrile (buffer B). For peptide separation, a linear gradient was employed beginning at 2% B, followed by a linear increase to 31% B over 17 minutes at a flow rate of 0.500 μl/min, then increased to 90% B for column wash before re-equilibration to initial conditions. The total runtime was 20 minutes per sample. Mass spectrometric analysis was performed in data-independent acquisition (DIA) mode with the ion source operated in positive polarity with a spray voltage of 1900 V, ion transfer tube temperature of 280°C, and RF lens value set to 40%. Full MS scans were acquired in the Orbitrap analyzer at a resolution of 240,000 over a scan range of 380-980 m/z with custom AGC target settings. Liquid chromatography mode was selected with an expected LC peak width of 10 seconds, and advanced peak determination was enabled. The default charge state was set to 2, and Orbitrap lock mass correction was turned off. For DIA analysis, the precursor mass range was set to 380-980 m/z, divided into 300 scan events with an isolation window of 2 m/z and window placement optimization enabled. MS2 spectra were acquired using HCD with a normalized collision energy of 25%. Fragment ions were detected using the Astral detector with a scan range of 150-2000 m/z. The normalized AGC target was set to 500% (absolute AGC value of 5.00e4) with a maximum injection time of 3 ms. Data were collected in centroid mode for MS2 scans with positive polarity. Quality control samples were analyzed regularly throughout the analytical sequence to monitor system performance and stability.

### Statistical Rationale

#### Data analysis / Spectral search

Raw MS files were converted to mzML format using ThermoRawFileParser (v4.3, Thermo Fisher Scientific) with format and metadata parameters set to 1 and 0, respectively. File conversion was performed in parallel on a high-performance computing cluster for efficient processing.

Spectral searches were conducted using DIA-NN (version 2.0) ^38^ against the UniProt human Swiss-Prot reference proteome database, including isoforms (version 19.02.2025). An in-silico spectral library was generated from the FASTA sequences incorporating oxidation of methionine and N-terminal acetylation as variable modifications. Trypsin specificity was set with up to 2 missed cleavages allowed. Mass accuracy tolerances were set to 10 ppm for both MS1 and MS2 levels, with a scan window parameter of 7.

Search parameters included match-between-runs functionality using the “--usequant” and “--reanalyse” flags, along with peak centering, smart profiling, retention time profiling, and relaxed protein inference settings. False discovery rate control was applied at 1% at the peptide-spectrum match level using a target-decoy approach. Protein quantification incorporated both MS1 and MS2 data without interference signal removal.

#### Bioinformatics analysis

The ‘Report.parquet’ output files from DIA-NN were used for all downstream analyses. All data processing and analysis steps were performed in the Python programming environment (v.3.12.10) using custom analysis scripts. Data processing was conducted using pandas (v2.2.3) for manipulation of proteomics data matrices and NumPy (v2.2.5) for numerical operations, while visualization and statistical analyses employed SciPy (v1.15.3), matplotlib (v3.10.3), and seaborn (v0.13.2). Although four technical replicates were measured for each column condition, only three replicates were included in the final analysis, with the first run of each column systematically excluded from the dataset. Visualization of raw files was conducted using the alpharaw package (https://github.com/MannLabs/alpharaw).

## Acknowledgement

We thank all members of the Proteomics and Signal Transduction Group for their help and discussions, in particular Igor Paron and Tim Heymann, for help with instrumentation. We further thank Johannes Maisch, Matthias Frübis, and Jan Schwerdt for the discussion and supply of test material that was used in this study.

## Author contribution

PEG and JBMR conceived and designed the study. ASS, KK, ECMI, and VA performed mass spectrometry experiments. ASS and KK carried out data processing and statistical analyses. ASS and KK interpreted results in the biological/clinical context. ASS, KK, and JBMR drafted the initial manuscript. All authors discussed results, revised the manuscript, and approved the final version.

## Supplementary Figures

**Supplementary Figure 1:**
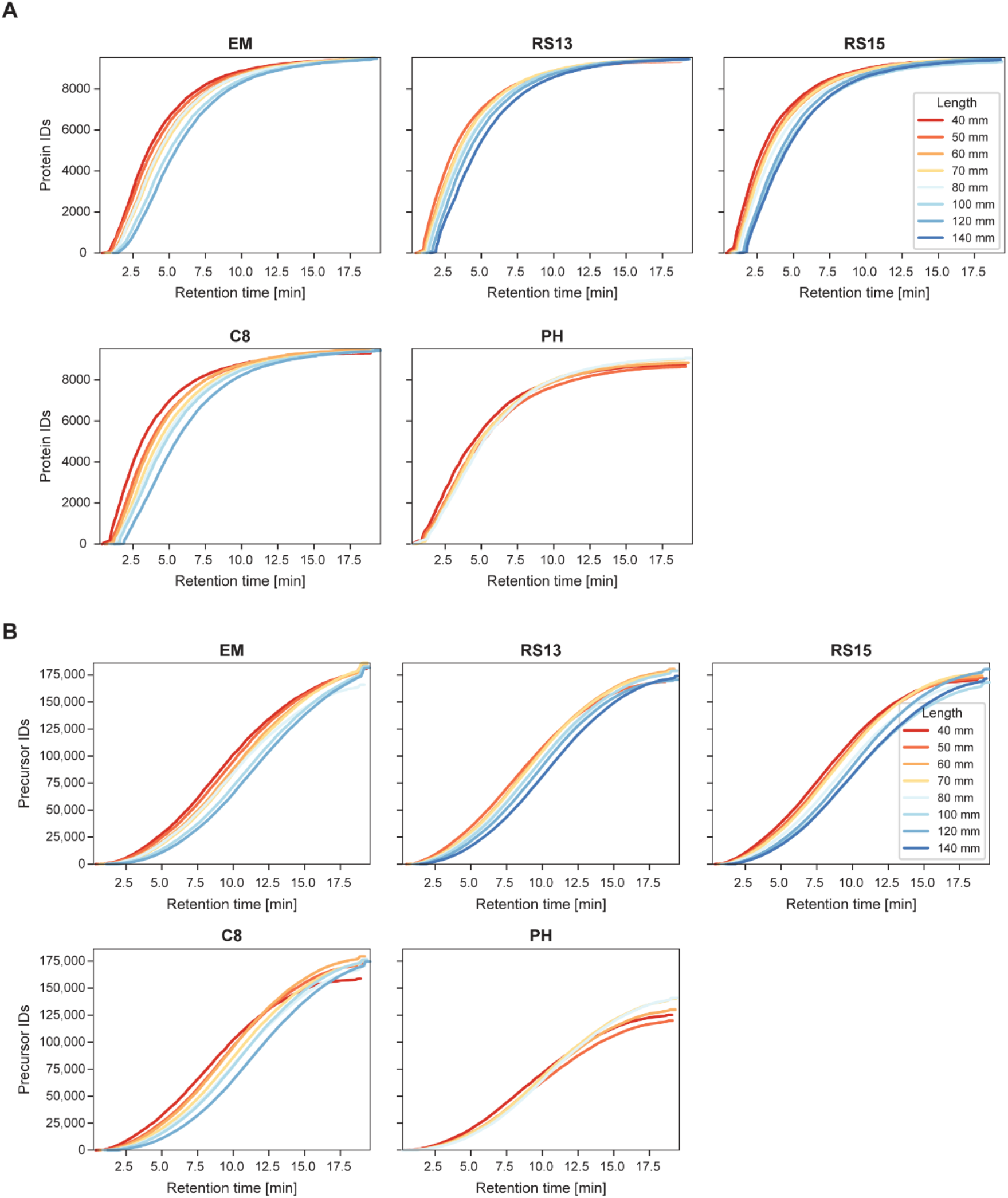
Cumulative identification profiles across column lengths. A: Cumulative protein identifications over retention time for each column chemistry. B: Cumulative precursor identifications over retention time for each column chemistry.

**Supplementary Figure 2:**
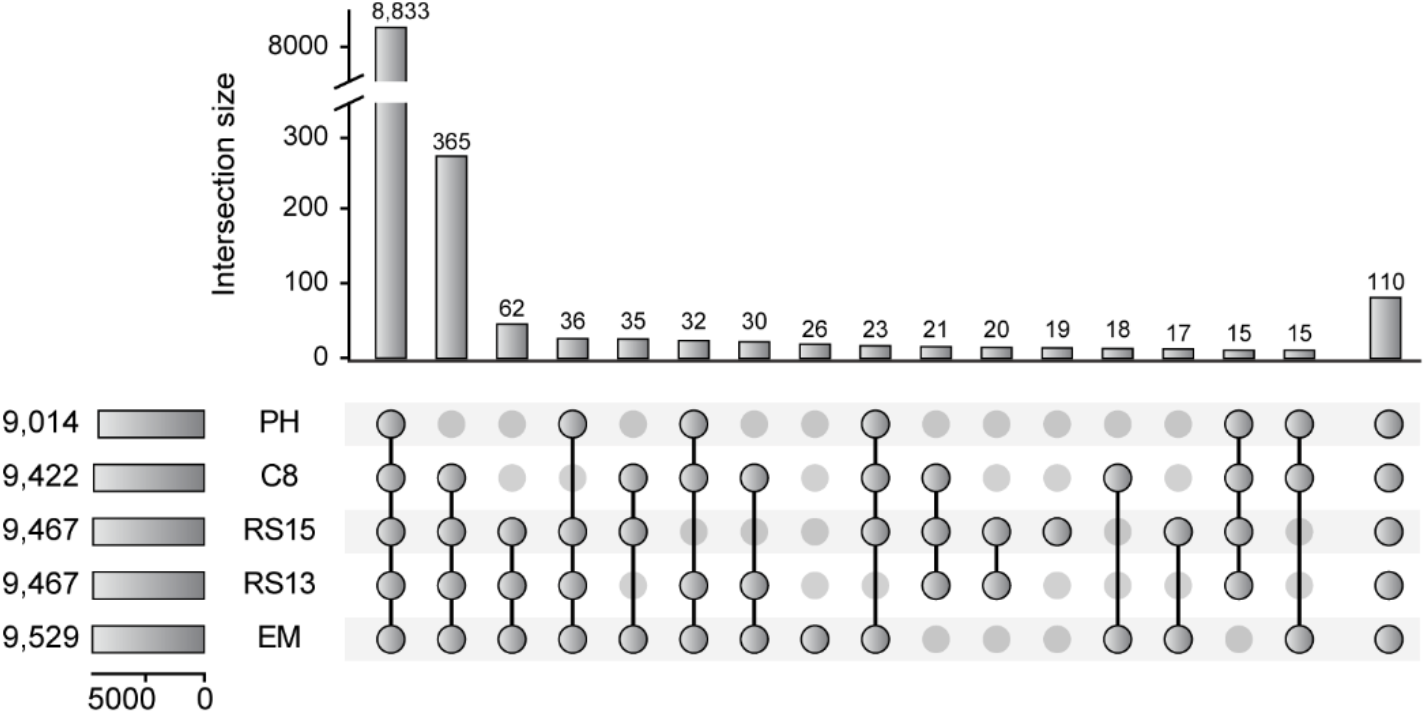
Protein group overlap across column chemistries at 70 mm. UpSet plot showing shared and unique protein groups between columns. Only the top intersections are shown; all remaining overlaps and unique sets are aggregated in the final bar.

**Supplementary Figure 3:**
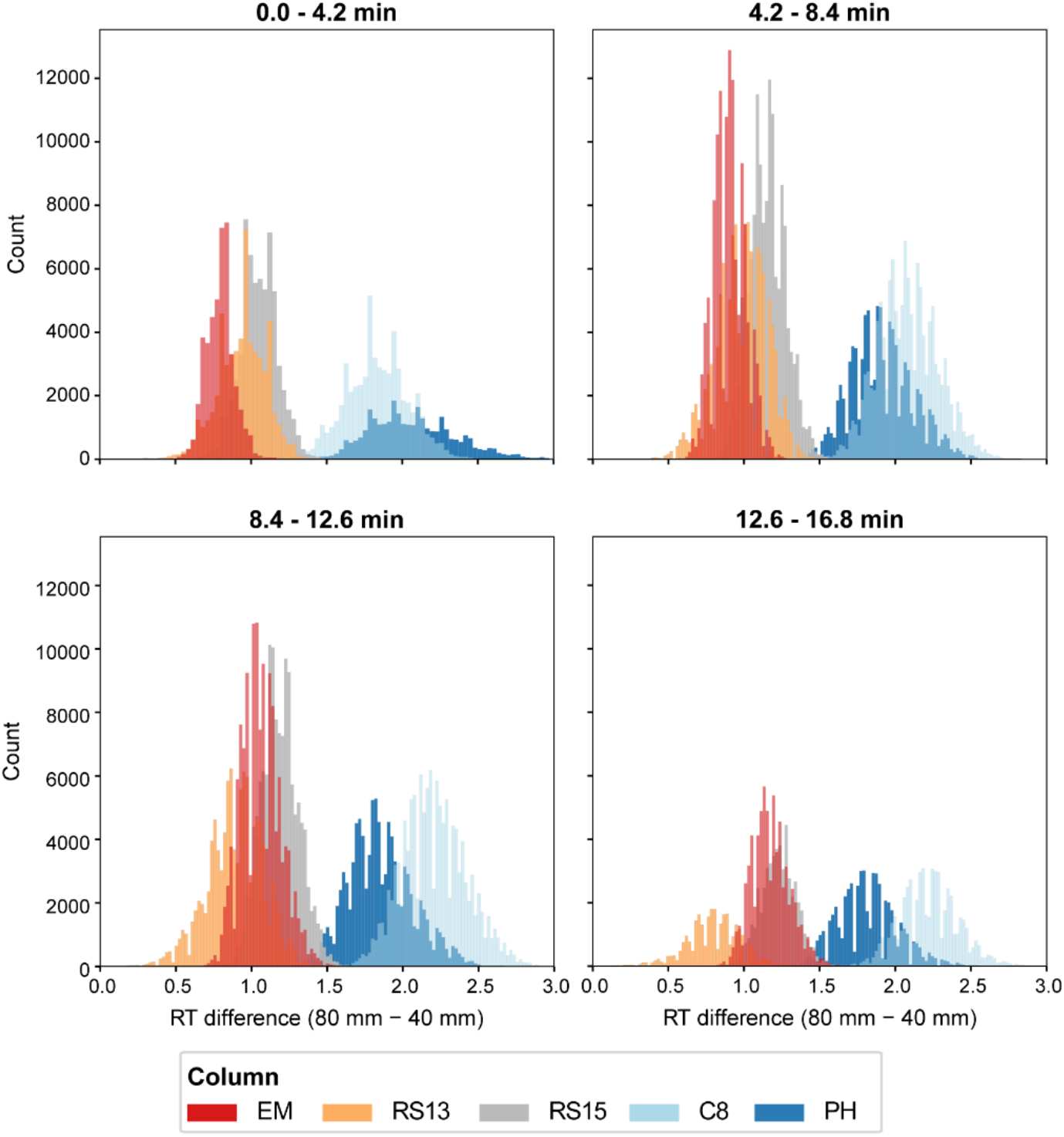
Retention time difference distributions between 80 mm and 40 mm columns stratified by elution time windows. Histograms show the distribution of retention time differences (ΔRT = RT_80_– RT_40_) for precursors binned into four retention time intervals: 0.0-4.2, 4.2-8.4, 8.4-12.6, and 12.6-16.8 minutes, based on their 40 mm column retention time. Each panel represents a specific elution window, with column chemistries color-coded as indicated in the legend. Precursors eluting after 16.8 minutes are not shown.

**Supplementary Figure 4:**
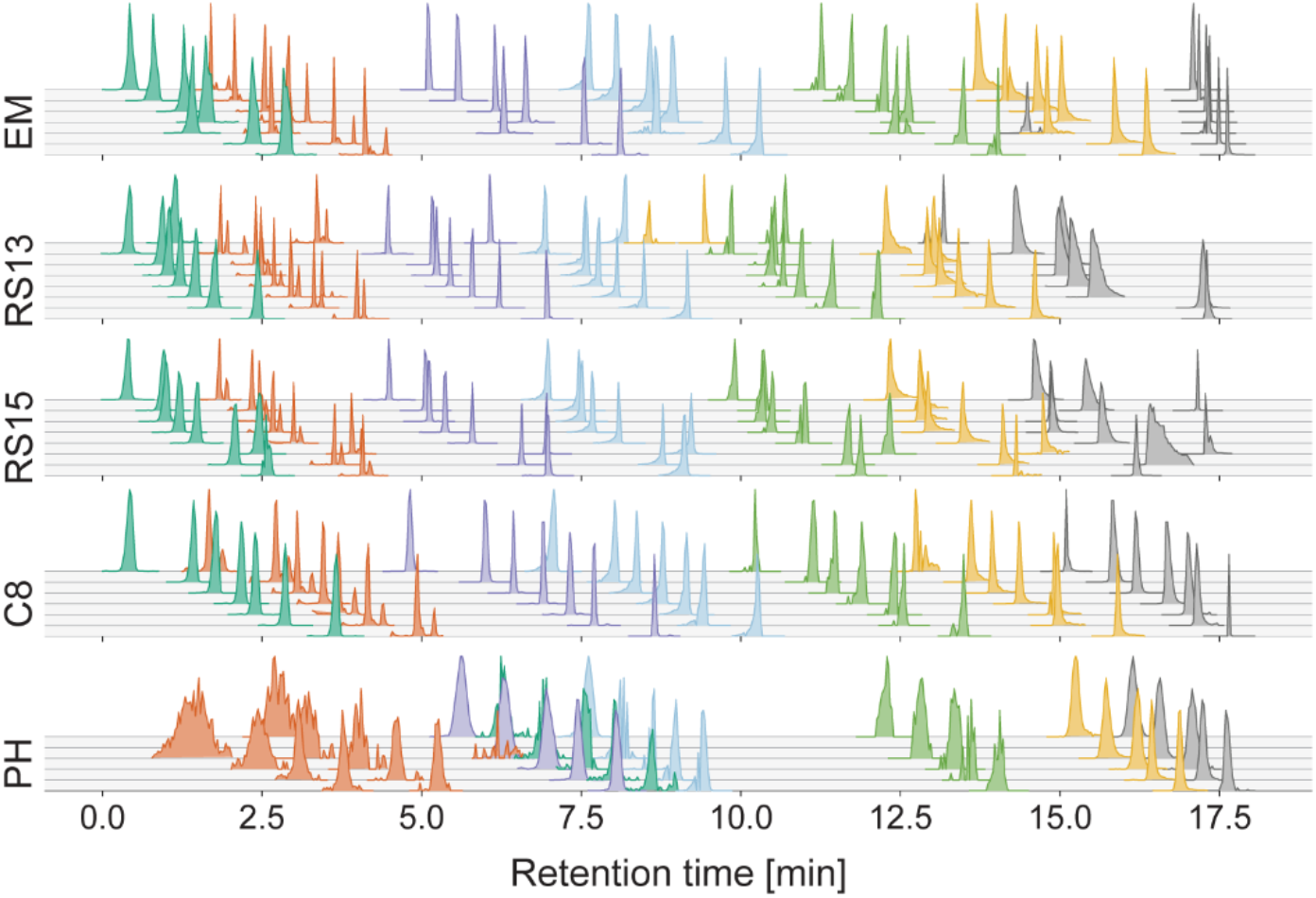
Extracted ion chromatograms of GAPDH precursors across column chemistries and lengths. Visualized replicate 3, selected precursors in retention time order: QASEGPLK2 (415.22433 m/z, dark green/turquoise), LTGMAFR2 (398.21274 m/z, orange), VGVNGFGR (403.2194 m/z, purple), GALQNIIPASTGAAK2 (706.3988 m/z, pink), LVINGNPITIFQERDPSK3 (681.0407 m/z, light green), WGDAGAEYVVESTGVFTTMEK3 (759.6842 m/z, yellow), VIHDNFGIVEGLMTTVHAITATQK4 (649.59546 m/z, grey). Precursors selected from >50 GAPDH precursors by filtering for intensity >4.0E+05 and retention time distribution. For each column material, chromatograms are arranged from top to bottom in order of increasing column length.

**Supplementary Figure 5:**
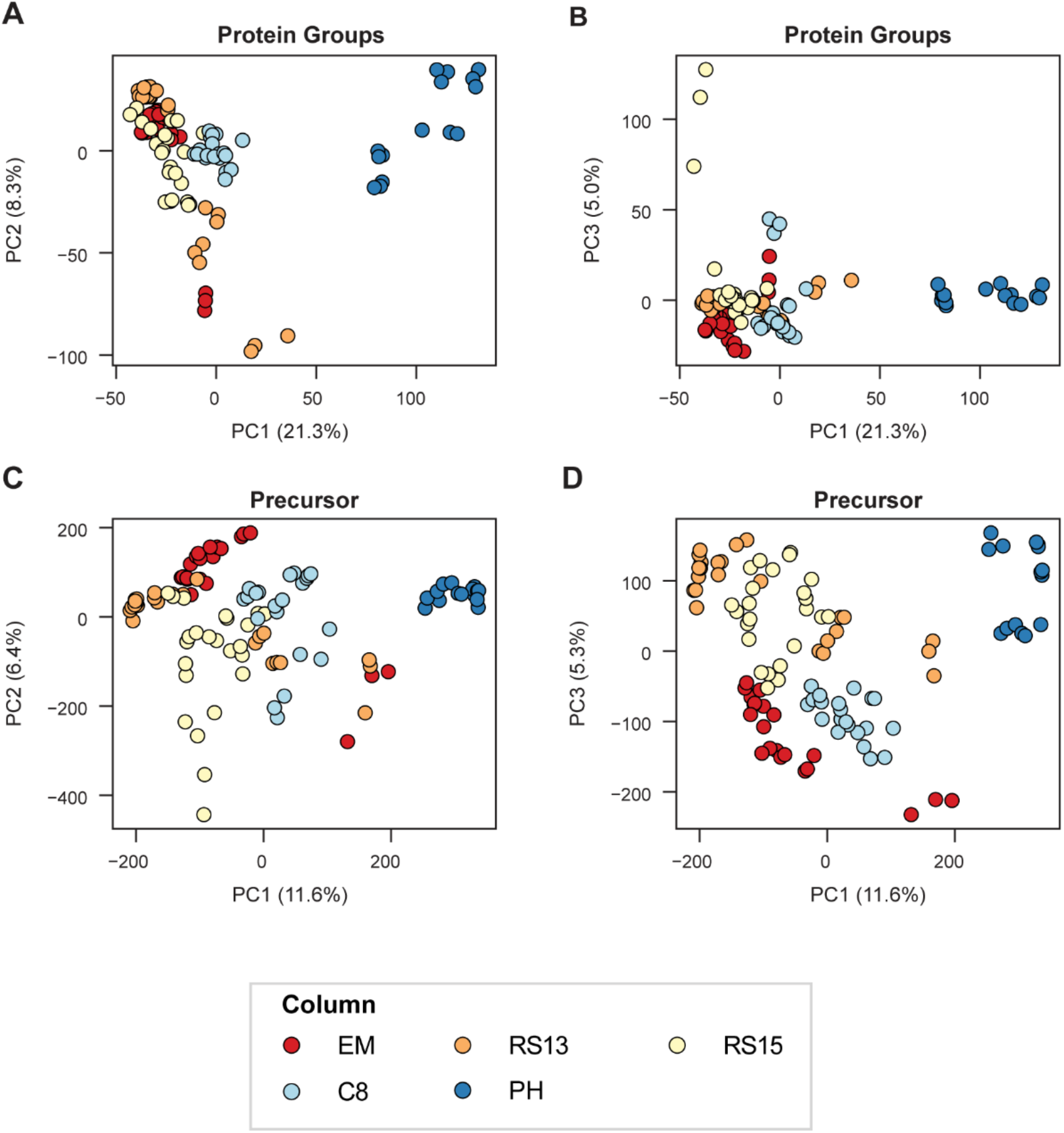
Principal component analysis (PCA) of identified protein groups and precursors. A, B: PCA of protein group intensities (PC1 vs. PC2 and PC1 vs. PC3). C, D: PCA of precursor intensities (PC1 vs. PC2 and PC1 vs. PC3). Panel C is identical to Figure 1E and is included here for completeness. All intensities were standardized (mean = 0, variance = 1) prior to PCA; missing values were imputed using k-nearest neighbor (k = 5).

**Supplementary Figure 6:**
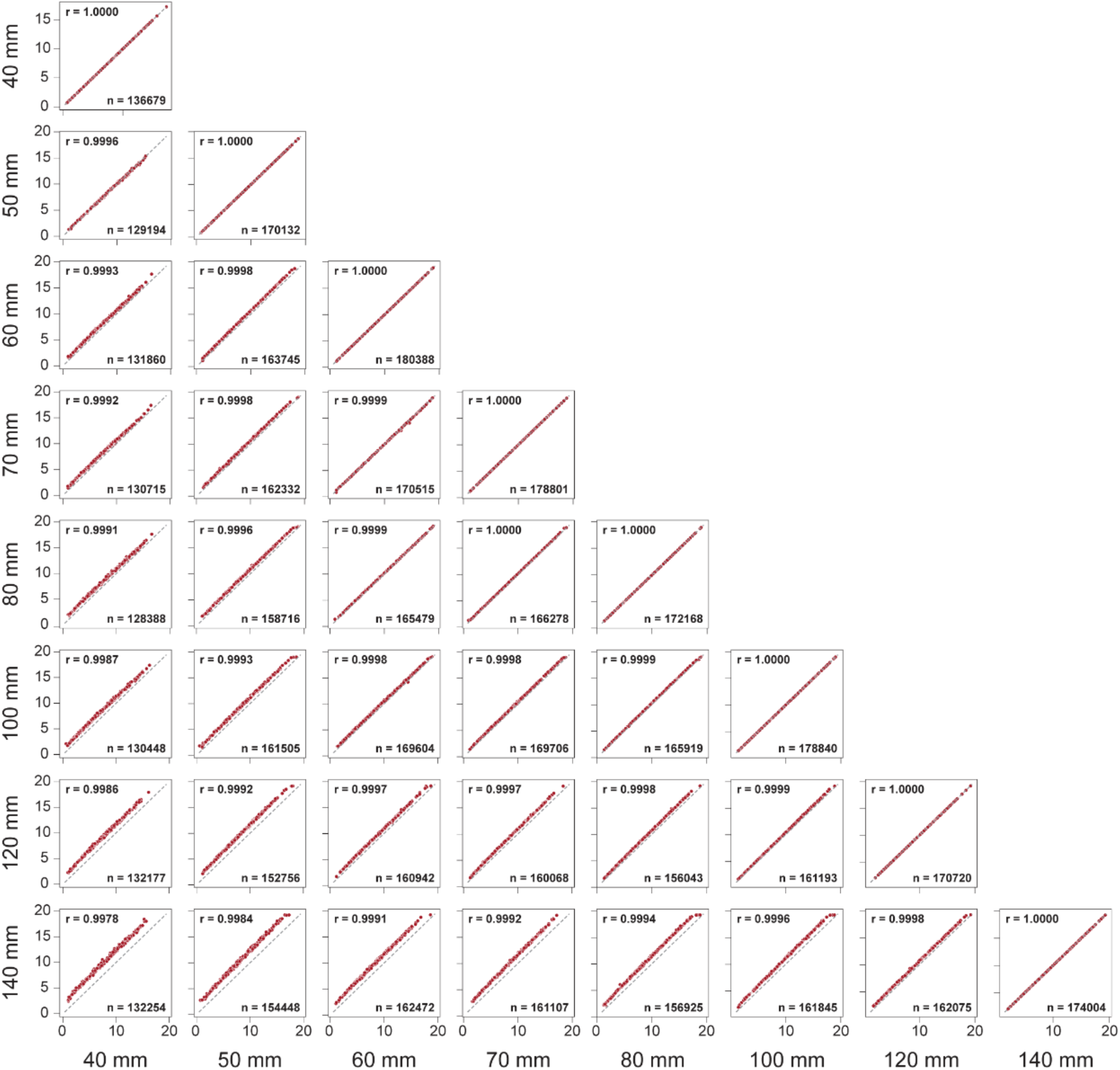
Retention time correlations across column lengths for RS13. Pairwise retention time correlations between all column lengths (40–140 mm) for RS13; only 0.2% of all data points are shown.

**Supplementary Figure 7:**
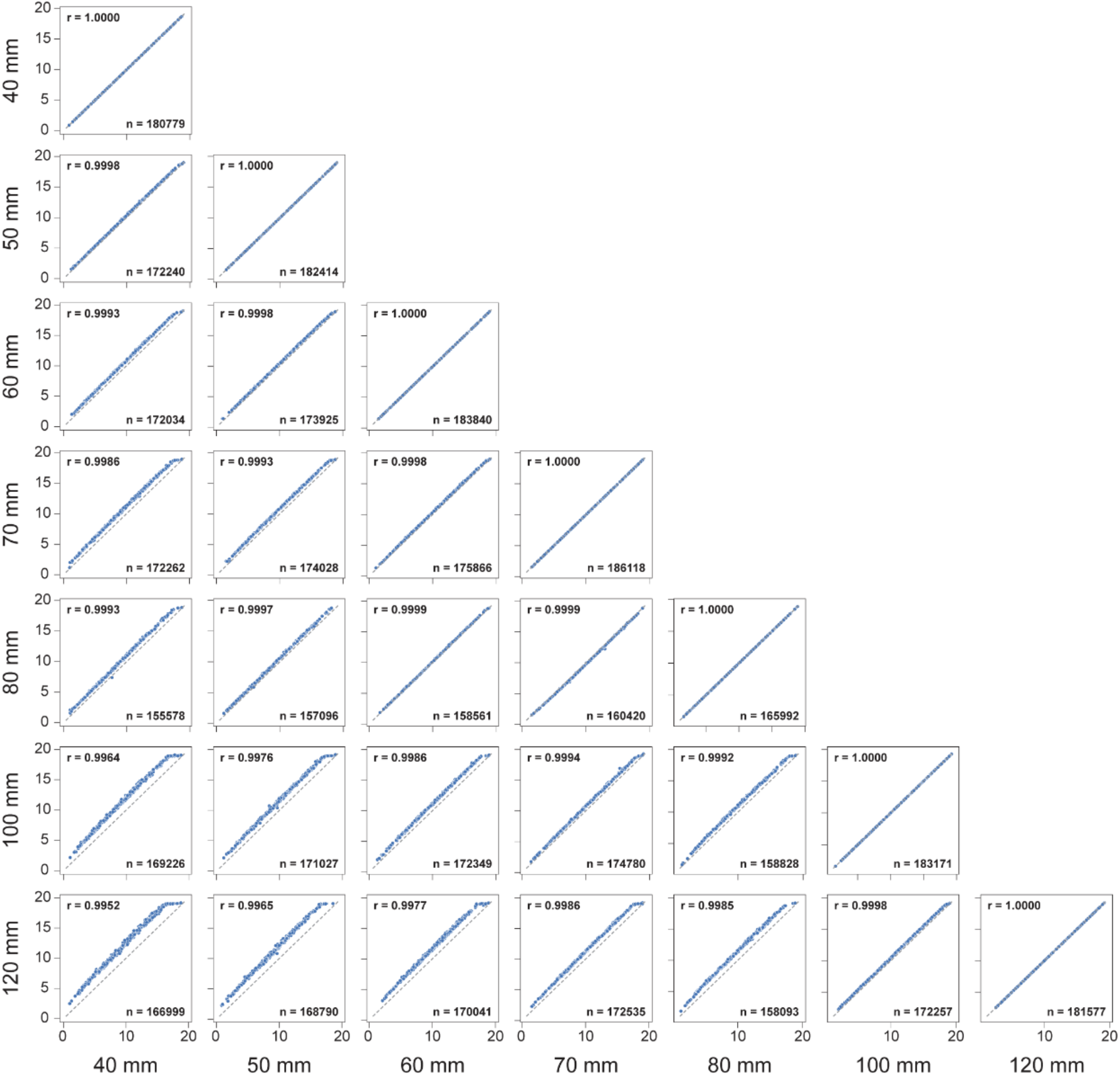
Retention time correlations across column lengths for EM. Pairwise retention time correlations between all column lengths (40–120 mm) for EM; only 0.2% of all data points are shown.

**Supplementary Figure 8:**
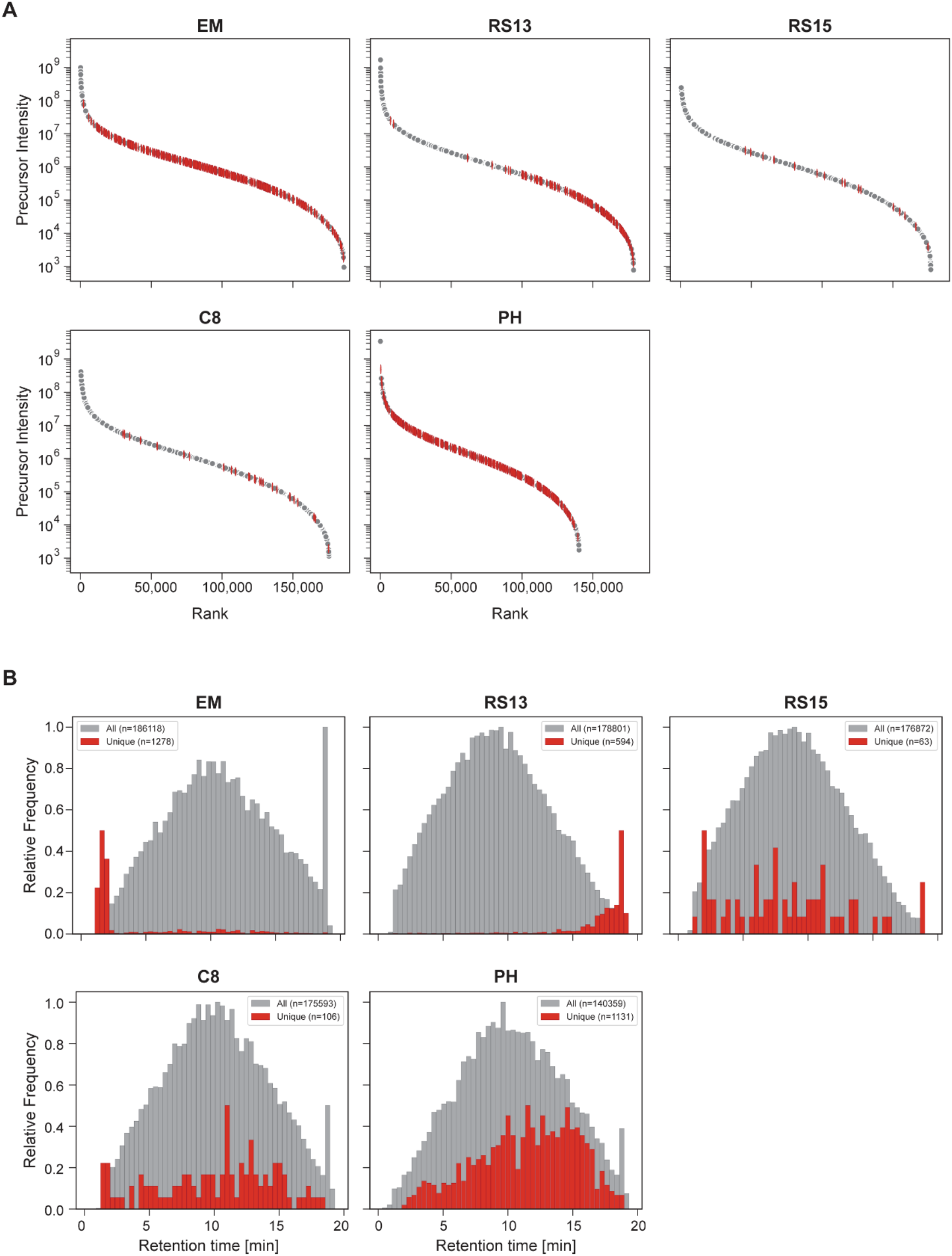
Unique precursor analysis for all column types. A: Rank abundance plots for each column showing all precursors (grey) and unique precursors (red). Only 1% of grey dots is shown and 25% of the red dots. B: Retention time distributions for each column. Distribution of all precursors is normalized to 1 and distributions of the unique precursors are normalized to 0.5.

